# TCTEX1D2 has two distinct functions in sperm flagellum formation in mice; cytoplasmic dynein 2 and inner dynein arm

**DOI:** 10.1101/2024.06.27.600786

**Authors:** Ryua Harima, Kenshiro Hara, Kentaro Tanemura

**Author notes:** **Competing interest:** The authors declare that no competing interests exist.

## Abstract

Flagella and cilia are widely conserved motile structures. Mammalian sperm possess flagella, which confer motility to the sperm and, therefore, are essential for fertilization. Large protein complexes called dynein, including cytoplasmic dynein 2 and axonemal dynein, play a role in the formation of cilia and flagella. The function of each dynein complex subunit in sperm flagellum formation is unclear. One such subunit is TCTEX1D2. *Tctex1d2*^−/−^ mice generated in this study showed male infertility because of flagellar dysplasia arising from round spermatids and disruption of the axonemal structure inside the flagella. In contrast, the motile cilia of *Tctex1d2*^−/−^ mice were normal. Co-immunoprecipitation studies showed that TCTEX1D2 interacted with the cytoplasmic dynein 2 component proteins WDR34, WDR60, and DYNLT1 in the testes and that the localization of these proteins was abnormal in *Tctex1d2*^−/−^ mice. Furthermore, TCTEX1D2 also interacted with WDR63 and WDR78, component proteins of the inner dynein arm, which is axonemal dynein. Overall, we revealed that TCTEX1D2 has two distinct functions in mouse sperm flagellum formation the assembly of cytoplasmic dynein 2 and organization of inner dynein arm which is not the case in cilia formation where TCTEX1D2 functions only as cytoplasmic dynein 2.

## Introduction

Flagella are widely conserved structures present in a large number of organisms ranging from protists to mammals, and their most distinctive feature is their motility ^1,2^. In mammals, the sperm is the only cell with flagella and thereby has motility, which is essential for fertilization ^3^. In addition to flagella, mammalian cells also have a similar motile structure called cilia ^4^. The flagellar and cilia axonemes have 9+2 microtubule internal structure, consisting of two central microtubules surrounded by nine peripheral microtubules ^5–7^. Large protein complexes called dynein, which move along the microtubules, play important roles in the formation of cilia and flagella and maintenance of motility ^8–10^. Dynein is composed of different subunits that are classified as heavy, intermediate, light intermediate, and light chain subunits based on their molecular weight ^11^. Dynein is divided into cytoplasmic and axonemal dynein, and cytoplasmic dynein is further divided into cytoplasmic dynein 1 and cytoplasmic dynein 2 ^12^. Cytoplasmic dynein 2 and axonemal dynein are closely related to cilia and flagella ^13–16^. Cytoplasmic dynein 2 primarily plays a role in cilia formation ^17,18^ and is responsible for retrograde transport as part of the intraflagellar transport (IFT) system within cilia ^18–20^. Axonemal dynein is present as part of the axonemal structure, and divided into outer dynein arm and inner dynein arm. Axonemal dynein generates a sliding force that allows the flagella and cilia to acquire motility ^15^. The functions of cytoplasmic dynein 2 and axonemal dynein are very different; therefore, most subunits are specific to either cytoplasmic or axonemal dynein ^21,22^. However, light chains are an exception to this rule. In *Chlamydomonas* flagella, some light chains are common between cytoplasmic dynein 2 and axonemal dynein ^23,24^. This suggests that dynein light chains are involved in many processes about related to flagella, ranging from formation to structural maintenance. In *Chlamydomonas*, dynein light chains contribute to the assembly of the components, structural stability, and motility of the flagella ^24^.

In mammals, most studies on dynein light chains have focused on somatic cilia, and very few have focused on sperm flagella. Thus, our understanding of the dynein light chains involved in sperm flagellum formation is very poor. Although flagella are widely conserved structures across species, it is not appropriate to apply the findings from other species to mammalian sperm flagella, because there are differences in the formation mechanisms and constituent proteins between species ^2,25–27^. The formation mechanism of sperm flagella and cilia is considered to be similar; however, sperm flagella are generally much longer than cilia and have unique structures, such as mitochondrial sheath, outer dense fiber, and fibrous sheath. Furthermore, it has been reported that the functions of proteins related to axoneme formation differ between sperm flagella and cilia ^16^, suggesting that there are differences in the mechanism of their formation. Therefore, it is important to use mammalian sperm flagella, to understand the function of dynein light chain in mammalian sperm flagella.

In the present study, we focused on Tctex1 domain-containing 2 (TCTEX1D2, also known as DYNLT2B) a dynein light chain present in the testes. In mammalian somatic cells, TCTEX1D2 is known as one of the cytoplasmic dynein 2 light chains and is involved in the formation of primary cilia (non-motile cilia) ^11,28,29^. Cytoplasmic dynein 2 has seven light chains (DYNLL1, DYNLL2, DYNLT1, DYNLT3, DYNLRB1, DYNLRB2, and TCTEX1D2) ^11,21^; however, the precise function of each subunit is unclear. Among the light chains, only TCTEX1D2 is specific to cytoplasmic dynein 2 ^11,28,29^, whereas the others are also found in cytoplasmic dynein 1 ^11^. In addition, DYNLL1 and DYNLL2, DYNLT1 and DYNLT3, and DYNLRB1 and DYNLRB2 are homologous to each other, and the homologs appear to have interchangeable functions ^30^. Therefore, TCTEX1D2 is unique and irreplaceable among the light chains of cytoplasmic dynein 2 and is thought to play a critical role in cytoplasmic dynein 2. However, there are no reports on the role of TCTEX1D2 in sperm flagellum formation, and it is yet unknown whether TCTEX1D2 is a component of cytoplasmic dynein 2 in germ cells. Interestingly, *Tctex1d2* mRNA is highly expressed in mouse testes according to NCBI gene (https://www.ncbi.nlm.nih.gov/gene/66061) and is mainly expressed in spermatocytes according to single-cell RNA-seq in mouse testes ^31^. Therefore, it is hypothesized that *Tctex1d2* might play a role in sperm flagellum formation. In this study, we aimed to elucidate the function of *Tctex1d2* in sperm flagellum formation.

In this study, we created *Tctex1d2*^−/−^ mice to investigate the function of *Tctex1d2*. Male *Tctex1d2*^−/−^ mice were infertile and displayed severe morphological abnormalities in sperm flagella and disorganized distribution of axonemal components. However, there was no effect on the formation of motile cilia; only the formation of sperm flagella was significantly impaired in *Tctex1d2*^−/−^ mice. TCTEX1D2 interacted with cytoplasmic dynein 2 component proteins in the testes, and we observed that the assembly of the cytoplasmic dynein 2 subunit is blocked in *Tctex1d2*^−/−^ mice. TCTEX1D2 also interacted with the subunits of axonemal dynein in the testes. This study revealed that TCTEX1D2 has two distinct functions cytoplasmic dynein 2 and axonemal dynein and plays an important role in sperm flagellum formation in mice.

## Results

### TCTEX1D2 is highly expressed in the testes and localized in flagella from round spermatids to elongated spermatids

*Tctex1d2* mRNA is highly expressed in mouse testes according to NCBI gene (https://www.ncbi.nlm.nih.gov/gene/66061). Western blotting (WB) was performed to investigate TCTEX1D2 protein expression. TCTEX1D2 was found to be highly expressed in the testes and spleen (fig. 1A). This shows that TCTEX1D2 may have certain functions in the testes. To determine the localization of TCTEX1D2 in germ cells, we performed immunohistochemistry (IHC) and immunocytochemistry (ICC) experiments. However, no antibodies were available for the IHC or ICC analyses. Therefore, we generated *Tctex1d2* - 3×FLAG mice, in which a 3×FLAG was knocked in at the N-terminus of *Tctex1d2* (fig. S1A) and implemented ICC in germ cells isolated from the testes of these mice. Spermatogenesis in *Tctex1d2* - 3×FLAG mice was comparable to that in WT mice, and the sperm flagella of *Tctex1d2* - 3×FLAG mice showed normal morphology (fig. S1B, C). Thus, 3×FLAG knock-in in *Tctex1d2* had no effect on the function of TCTEX1D2. We found that TCTEX1D2 was localized along the flagella from round spermatids to elongated spermatids. (fig. 1B-D). TCTEX1D2 was also localized in the manchette, which are transiently formed in step 10–14 spermatids (fig. 1C). Furthermore, TCTEX1D2 was also expressed in the cilia of ventricles, trachea, and oviducts (fig. S2A-C).

**Figure. 1.**
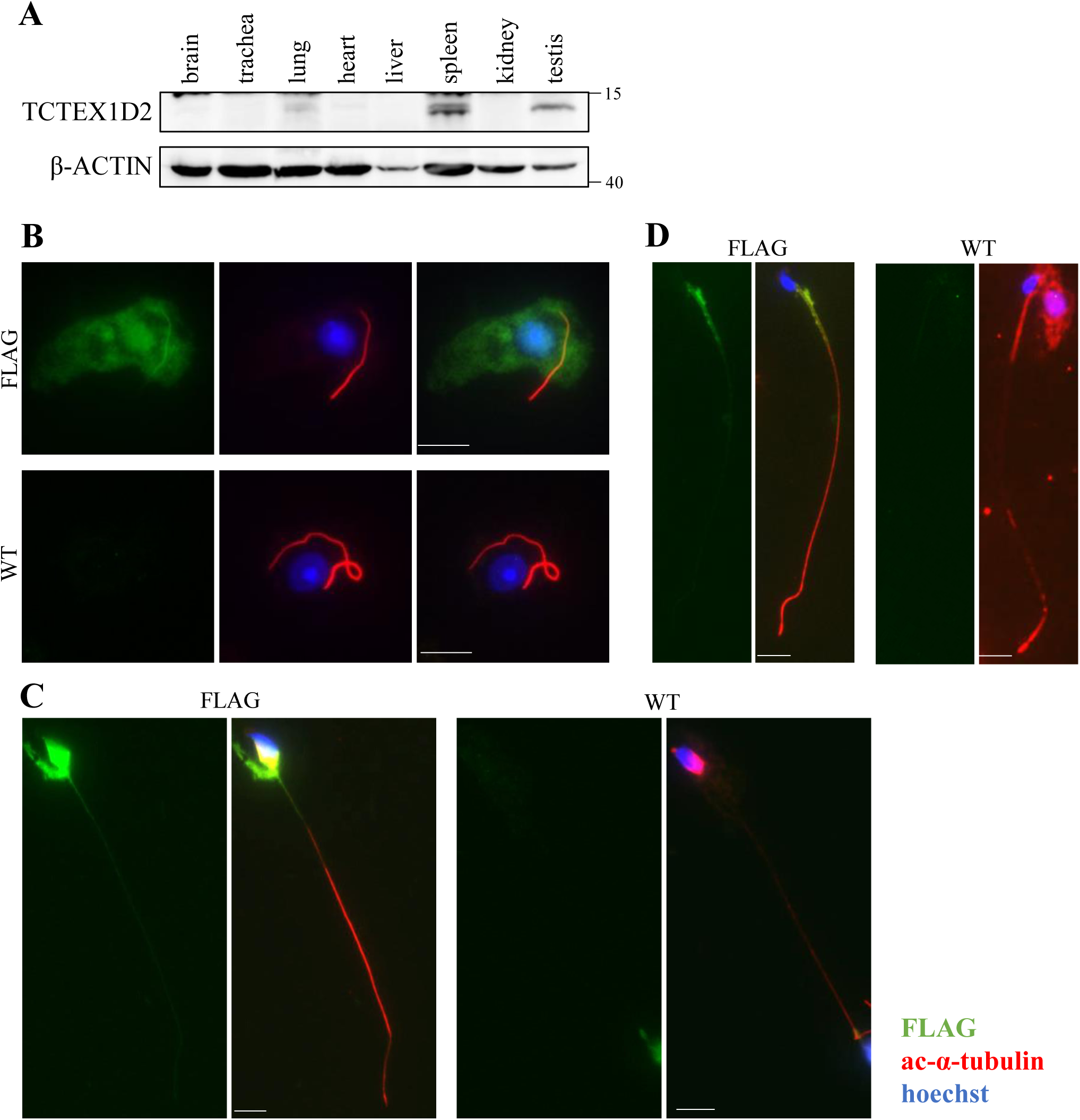
TCTEX1D2 was high expression in the testis and localized along the flagella from round spermatids to elongated spermatids and the manchette. (A) Protein expression analysis of mouse TCTEX1D2 in various organs using western blotting (WB); TCTEX1D2 was highly expressed in the testes and spleen. (B-D) ICC in WT and *Tctex1d2* - 3×FLAG germ cells to analyze the localization of TCTEX1D2 (green: anti-FLAG, red: anti-acetylated-α-tubulin, blue: hoechst33342). TCTEX1D2 localized to flagella from early round spermatids to elongated spermatids. In addition, TCTEX1D2 localized to the manchette. (B) Round spermatids in steps 1–3. (C) Elongated spermatids of steps 10–12. (D) Elongated spermatids of steps 15–16. Scale bars are 10 µm.

### TCTEX1D2 is a component of cytoplasmic dynein 2

In mammalian somatic cells, TCTEX1D2 is a component of cytoplasmic dynein 2 (fig. 2A). Therefore, we analyzed whether TCTEX1D2 is a component of cytoplasmic dynein 2 in germ cells. Cytoplasmic dynein 2 consists of 11 subunits: heavy chain (DYNC2H1), intermediate chain (WDR34 and WDR60), light intermediate chain (DYNC2LI1), and light chain (DYNLRB1, DYNLRB2, DYNLL1, DYNLL2, DYNLT1, DYNLT3, and TCTEX1D2) (fig. 2A). In mammalian somatic cells, TCTEX1D2 interacts with DYNLT1 and WDR60 ^30^. In the testes, TCTEX1D2 interacted not only with WDR60 and DYNLT1, but also with WDR34 (fig. 2B). These results indicate that TCTEX1D2 is one of the components of cytoplasmic dynein 2 in germ cells.

**Figure. 2.**
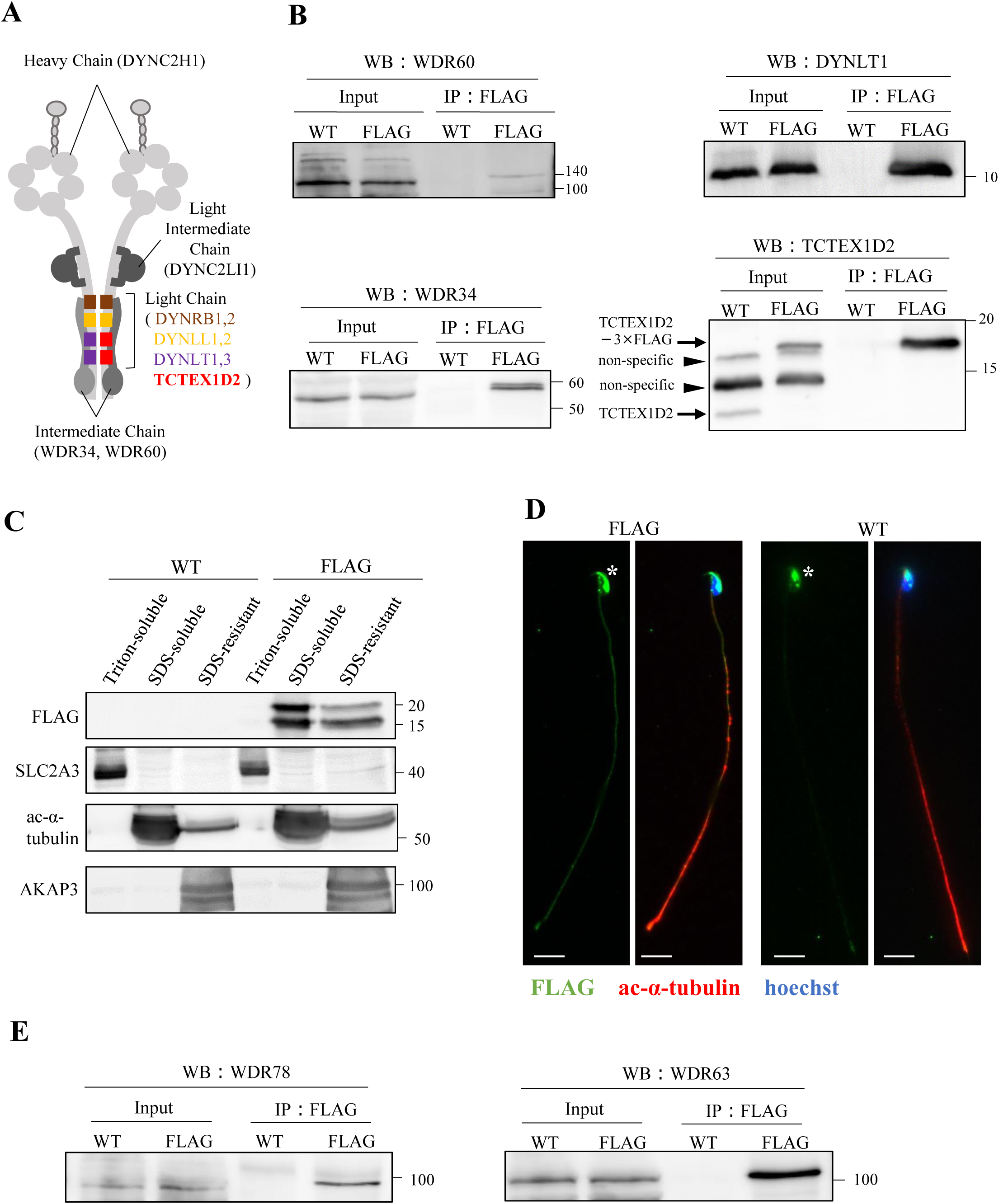
TCTEX1D2 interacted with the subunits of cytoplasmic dynein 2 and axonemal dynein. (A) Schematic diagram of the cytoplasmic dynein 2 complex in somatic cell. (B) Co-immunoprecipitation in the testes of *Tctex1d2* - 3×FLAG mice. In the testes, TCTEX1D2 interacted with WDR60, WDR34, and DYNLT1. (C) WB of cauda epididymis sperm fractionated into Triton X-100 soluble, SDS-soluble, and SDS-insoluble fractions from WT and *Tctex1d2* - 3×FLAG mice. TCTEX1D2 detected in SDS-soluble and SDS-insoluble fractions. (D) ICC of cauda epididymis sperm from WT and *Tctex1d2* - 3×FLAG mice (green: FLAG, red: acetylated-α-tubulin, blue: hoechst33342). TCTEX1D2 localized along the flagella. Asterisks indicate non-specific signals. Scale bars are 10 µm. (E) Co-immunoprecipitation with WDR63 and WDR78 in the testes of *Tctex1d2* - 3×FLAG mice. TCTEX1D2 interacted with WDR63 and WDR78. The used membrane was same as that in (B).

### TCTEX1D2 is an axoneme component and interacts with subunits of the inner dynein arm

In the flagella of *Chlamydomonas*, Tctex2b (a homolog of mouse TCTEX1D2) is present as a light chain of the inner dynein arm in the axoneme dynein ^32^. The inner dynein arm is important for controlling the size and shape of the bend of the flagella, features collectively referred to as waveforms ^33^. In *Chlamydomonas*, loss of function of the inner arm dynein subunit causes impaired flagellar motility ^34–36^. Thus, it is believed that in mammals, TCTEX1D2 is present in the sperm flagella as a light chain of the inner dynein arm. We analyzed the protein expression of TCTEX1D2 in the spermatozoa of the cauda epididymis. WB analysis of sperm proteins fractionated into Triton X-100 soluble, SDS soluble, and SDS-insoluble fractions was performed. SLC2A3, Ac-α-tubulin, and AKAP3 are markers for sperm membrane, axoneme, and other insoluble fractions, respectively. TCTEX1D2 was present in both the SDS-soluble and SDS-resistant fractions (fig. 2C). ICC studies also revealed that TCTEX1D2 localized along the sperm flagella (fig. 2D). These results showed that TCTEX1D2 is an axoneme component in mice. Further, co-immunoprecipitation studies revealed that TCTEX1D2 interacted with WDR63 and WDR78 (fig. 2E). Both proteins are intermediate chains of the inner dynein arm, which is a component of the 9+2 microtubule structure. These results indicate that TCTEX1D2 has some functions in the light chain of the inner dynein arm in mouse sperm flagella, as well as in that of *Chlamydomonas* flagella.

### *Tctex1d2* is essential for male fertility

To explore the function of TCTEX1D2 in the mouse testes, we generated *Tctex1d2* knockout (*Tctex1d2*^−/−^) mice using the CRISPR/Cas9 system (fig. 3A). We confirmed a 4696-bp deletion in the generated knockout mice using genotyping and Sanger sequencing (fig. 3B, C). Furthermore, the generated knockout mice completely lost TCTEX1D2 protein expression, as shown by results of WB experiments (fig. 3D). Fertility tests were conducted to assess the fertility of *Tctex1d2*^−/−^ male mice. *Tctex1d2*^−/−^ male mice were mated with two wild-type (WT) female mice. *Tctex1d2*^−/−^ male mice were found to be completely infertile (fig. 3E). In contrast, the testis/body weight ratio was unchanged in *Tctex1d2*^−/−^ mice in comparison with *Tctex1d2*^+/+^ mice (fig. 3F).

**Figure. 3.**
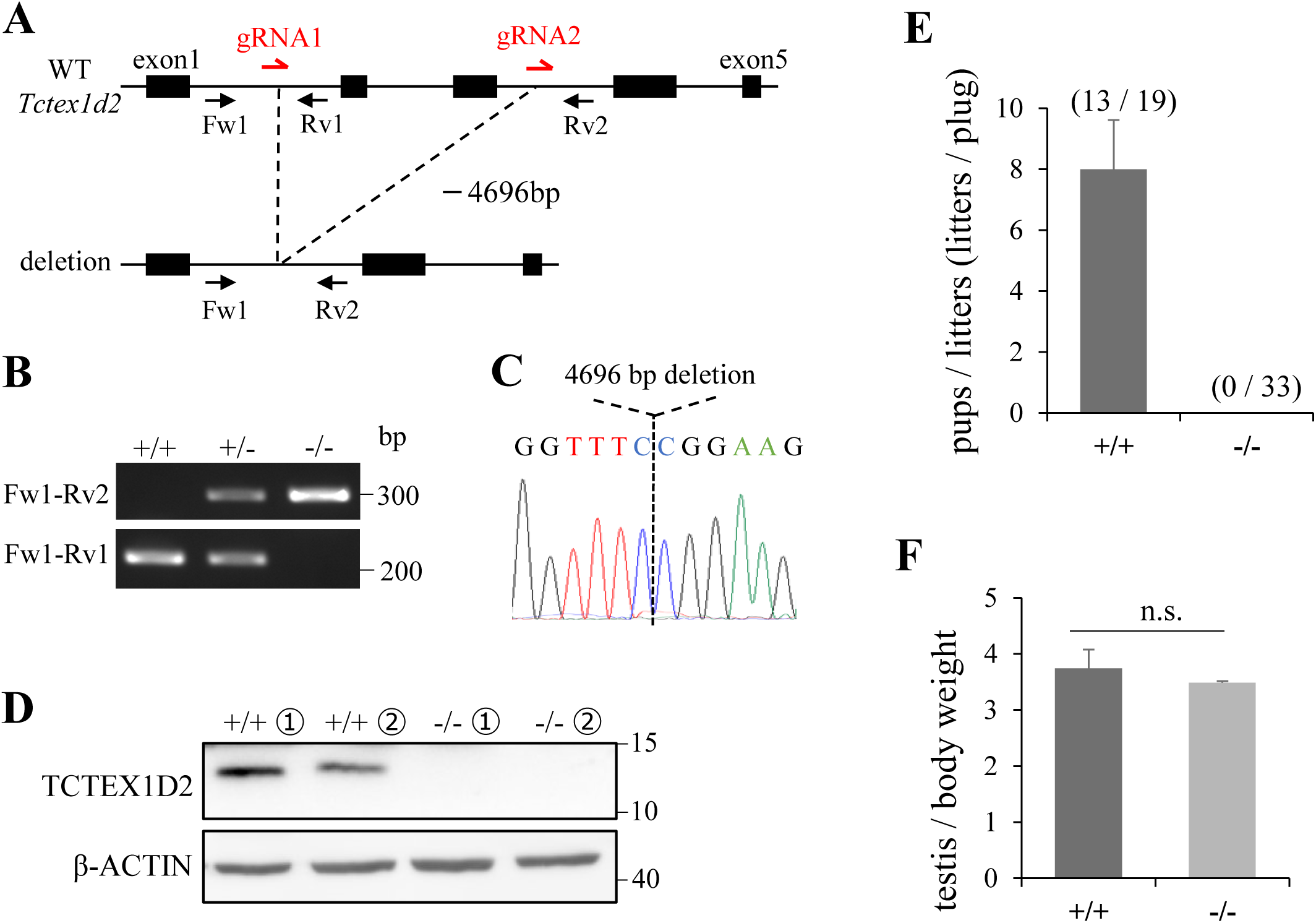
TCTEX1D2 was required for male fertility. (A) Schematic representation of the generation of *Tctex1d2*^−/−^ mice. The gRNA was designed between exons 1 and 2 and between exons 3 and 4. Genotyping primer pairs were Fw1 and Rv1 for *Tctex1d2*^+/+^ mice, Fw1 and Rv2 for *Tctex1d2*^−/−^ mice. (B) Genotyping and (C) Sanger sequencing showed that *Tctex1d2*^−/−^ mice had a 4696 bp deletion within the target region. (D) WB of tissue from mouse testes confirmed that TCTEX1D2 protein expression was completely lost in *Tctex1d2*^−/−^ mice. (E) Male *Tctex1d2*^+/+^ and *Tctex1d2*^−/−^ mice were mated with two female WT mice. We observed 33 plugs in *Tctex1d2*^−/−^ mice; however, *Tctex1d2*^−/−^ mice were unable to sire any offspring. (F) The ratio of testes weight / body weight. There were no differences between *Tctex1d2*^+/+^ and *Tctex1d2*^−/−^ mice. Error bars indicate means ± S.E. Two-tailed Student’s t-test, n=3, n.s. : no significant.

### *Tctex1d2*^−/−^ mice exhibit abnormal morphology of sperm flagella, but not of cilia

To identify the cause of infertility in *Tctex1d2*^−/−^ male mice, we performed periodic acid-Schiff hematoxylin (PAS-H) staining of the testes and epididymis. In stage I-III seminiferous tubules, a layered structure of germ cells was observed (fig. 4A, yellow box) and numerous spermatid flagella were present in the tubule lumen of *Tctex1d2*^+/+^ mice (fig. 4A, red box arrow). In contrast, a layered structure of germ cells was observed (fig. 4B, yellow box), but only a few flagella were present in the tubule lumen of *Tctex1d2*^−/−^ mice (fig. 4B, red box). In stage Ⅶ seminiferous tubules, a large number of elongated spermatids were observed in the lumen in *Tctex1d2*^+/+^ mice (fig. 4C). However, only a few elongated spermatids were observed in the lumen of *Tctex1d2*^−/−^ mice (fig. 4D), and the number of spermatozoa was significantly decreased in the cauda epididymis (fig. 4E, F). Furthermore, almost all the sperm flagella of *Tctex1d2*^−/−^ mice showed markedly abnormal morphology compared with those of *Tctex1d2*^+/+^ mice (fig. 4G), such as shortened and coiled tails (fig. 4H). In mice, the epithelial cells of the ventricles, trachea, and oviducts have motile cilia similar to sperm flagella. We analyzed the morphology of these cilia using Hematoxylin-Eosin (HE) staining. In contrast to the severe defects in sperm flagella, the cilia of the ventricles, trachea, and oviducts showed no abnormal morphology in *Tctex1d2*^−/−^ mice (fig. S3A–C). Therefore, we concluded that only the sperm flagella showed severely abnormal morphology in *Tctex1d2*^−/−^ mice.

**Figure. 4.**
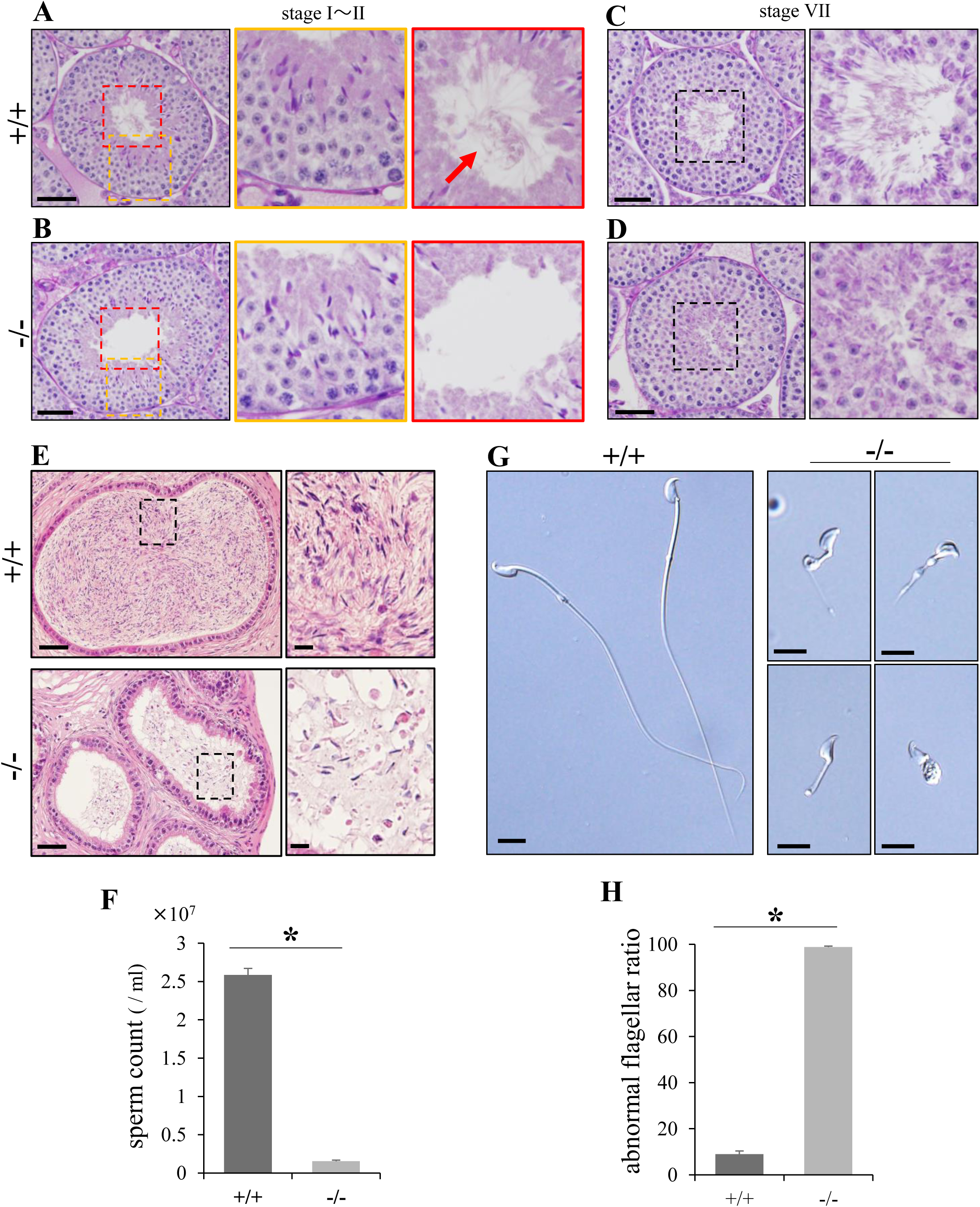
*Tctex1d2* is important for sperm flagellum formation. (A–D) Periodic acid Schiff hematoxylin (PAS-H) stain of seminiferous tubules of *Tctex1d2*^+/+^ and *Tctex1d2*^−/−^ mice. In stage I–II seminiferous tubules, *Tctex1d2*^−/−^ mice showed layered structures of germ cells (B; yellow box), similar to that in *Tctex1d2*^+/+^ mice (A; yellow box). However, sperm flagella were barely observed in the lumen in *Tctex1d2*^−/−^ mice (B; red box) compared to *Tctex1d2*^+/+^ mice (A; red box arrow). In the lumen of stage VII seminiferous tubules, only a few elongated spermatids were observed in *Tctex1d2*^−/−^ mice (D; black box) in contrast to *Tctex1d2*^+/+^ mice (C; black box). Scale bars are 50 µm. (E) Hematoxylin and eosin (H&E) stain of the cauda epididymis. Many spermatozoa were observed in *Tctex1d2*^+/+^ mice, whereas only very few spermatozoa were observed in *Tctex1d2*^−/−^ mice. Scale bars are 50 µm. (F) Sperm counts in the cauda epididymis. *Tctex1d2*^−/−^ mice had significantly reduced sperm counts compared to *Tctex1d2*^+/+^ mice. Error bars are means ± S.E. Two-tailed Student’s t-test, n=3, \**p* <0.05. (G) Cauda epididymis sperm observed using phase contrast microscopy. Sperm flagella of *Tctex1d2*^−/−^ mice showed morphological abnormalities. Scale bars are 10 µm. (H) The ratio of sperm with abnormal flagella in the cauda epididymis. Almost all sperm showed abnormal flagella in *Tctex1d2*^−/−^ mice. Error bars are means ± S.E. Two-tailed Student’s t-test, n=3, \**p* <0.05.

### *Tctex1d2*^−/−^ mice show abnormal sperm flagella and disruption of axoneme structures in early round spermatids

The results shown in fig. 1B-D indicate that TCTEX1D2 is expressed from the early stage of sperm flagellum formation and may have some functions at that stage. To further understand the role of TCTEX1D2, we analyzed flagellum formation during spermiogenesis. Flagellum formation in *Tctex1d2*^−/−^ mice was normal until steps 2–3 of the spermatids (fig. 5A). However, the axoneme failed to elongate normally and showed significant defects from step 4 spermatids onwards in *Tctex1d2*^−/−^ mice (fig. 5A). Using transmission electron microscopy, we examined the internal structure of the flagella and found that the axonemal components were disorganized in *Tctex1d2*^−/−^ mice. Doublet microtubules were not formed and many single microtubules were observed in *Tctex1d2*^−/−^ mice (fig. 5B). Moreover, the 9+2 microtubule structure, inner dynein arm (IDA), outer dynein arm (ODA), outer dense fiber (ODF), and fibrous sheath (FS) were disorganized and scattered in *Tctex1d2*^−/−^ mice (fig. 5B). Thus, the cause of the failure of flagellar elongation in *Tctex1d2*^−/−^ mice was disruption of the axonemal structure. These results suggested that TCTEX1D2 plays an important role in the organization of axonemal components.

**Figure. 5.**
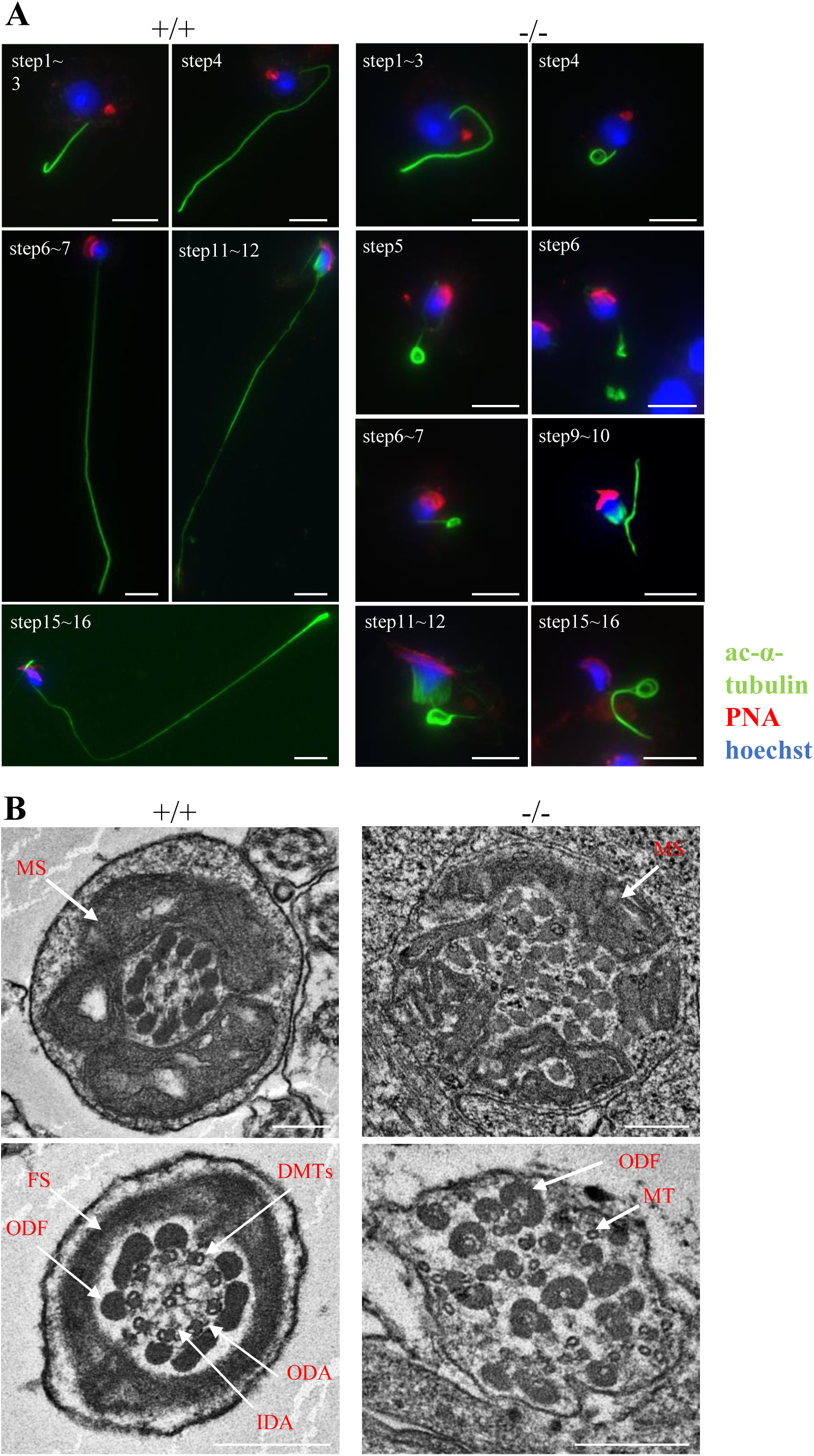
Abnormal assembly of sperm flagella in *Tctex1d2*^−/−^ mice occurred from step 4 round spermatids. (A) Flagella formation in spermiogenesis using ICCs (green: anti-acetylated-α-tubulin, red: PNA lectin, blue: hoechst33342). In *Tctex1d2*^+/+^ mice, flagella elongated as the spermatids differentiated. However, *Tctex1d2*^−/−^ mice showed markedly abnormal flagella after step 4 spermatids. Scale bars are 10 µm. (B) Observation of the internal structure of the flagella of spermatids using transmission electron microscopy. In *Tctex1d2*^+/+^ mice, microtubule 9+2 structures and other structures were neatly aligned in the axonemes. In contrast, in *Tctex1d2*^−/−^ mice, the internal structure of the axonemes were disorganized. The 9+2 structures, inner dynein arm (IDA), outer dynein arm (ODA), outer dense fiber (ODF), and fibrous sheath (FS) were not aligned properly. Scale bars are 250 nm. MS, mitochondrial sheath; DMTs, doublet microtubules; ODF, outer dense fibers; FS, fibrous sheath; ODA, outer dynein arm; IDA, inner dynein arm; MTs, microtubule (not doublet).

### TCTEX1D2 functions in the assembly of cytoplasmic dynein 2 subunits

TCTEX1D2 is one of the components of cytoplasmic dynein 2 in germ cells. Therefore, we investigated the expression patterns of TCTEX1D2 interacted subunit (WDR60, WDR34, and DYNLT1) in *Tctex1d2*^−/−^ mice using ICC. WDR60 and WDR34 were not expressed in the flagella both *Tctex1d2*^+/+^ and *Tctex1d2*^−/−^ mice round spermatids (fig. 6A, B). In the flagella in *Tctex1d2*^+/+^ mice elongated spermatids, WDR60 localized along the flagellar, especially strongly in principal piece (fig. 6A). WDR34 also localized along the flagellar, especially strongly in middle piece (fig. 6B). However, expression patterns of WDR60 and WDR34 were abnormal in *Tctex1d2*^−/−^ mice elongated spermatids, including loss of signal (fig. 6A, B above) and ectopic signal (fig, 6A, B bellow arrow head). DYNLT1 uniformly localized along the flagellar in *Tctex1d2*^+/+^ mice round spermatids to elongated spermatids (fig. 6C). However, in *Tctex1d2*^−/−^ mice, expression of DYNLT1 decreased in the flagella of round spermatids and loss in elongated spermatids (fig. 6C). These results suggest that TCTEX1D2 plays a role in the assembly and structural stability of the cytoplasmic dynein 2 subunits.

**Figure. 6.**
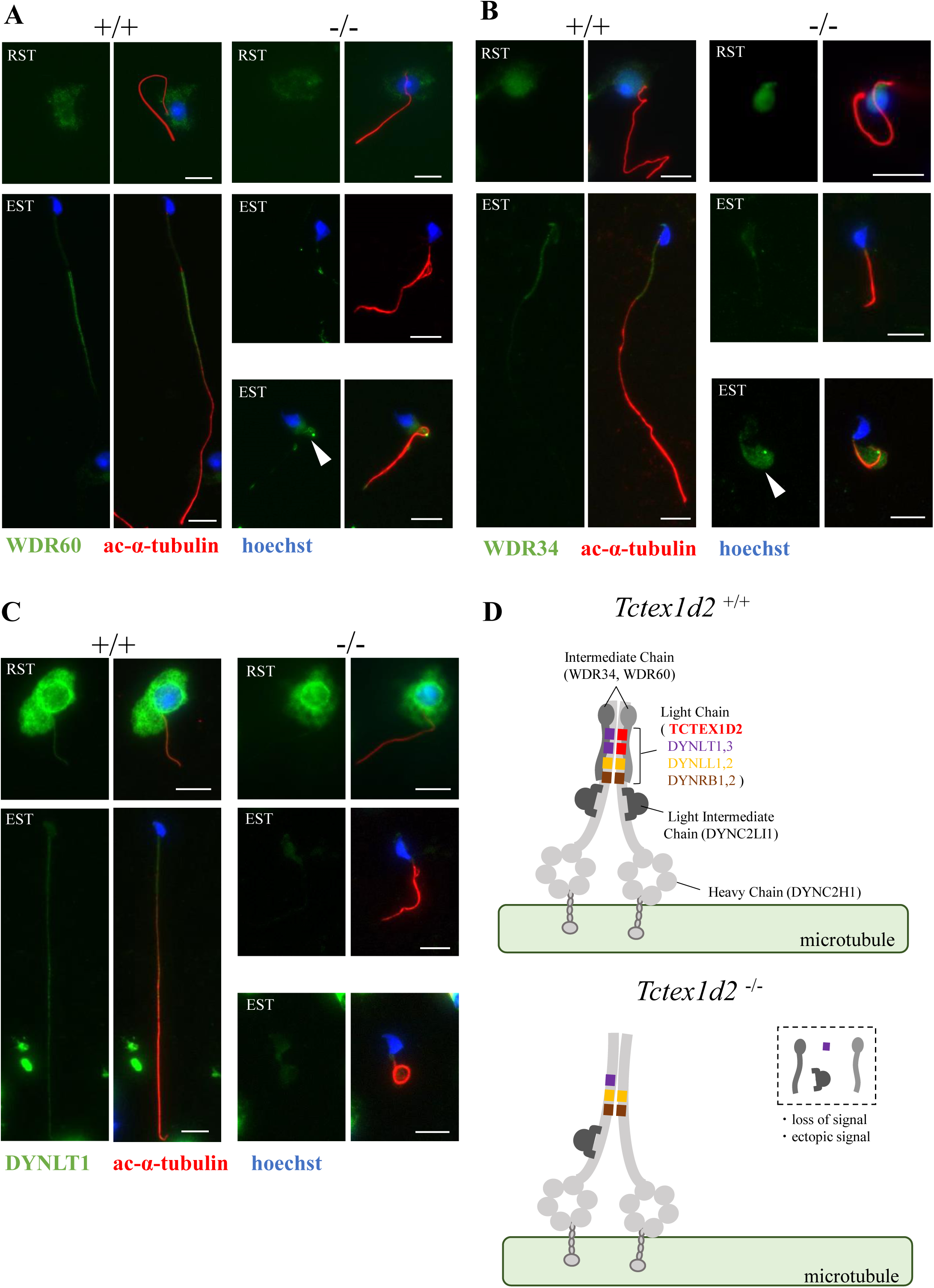
TCTEX1D2 played a role in the assembly of cytoplasmic dynein 2 subunits. (A) Expression analysis of WDR60 in *Tctex1d2*^−/−^ mice using ICC (green: WDR60, red: acetylated-α-tubulin, blue: hoechst33342). There was no expression of WDR60 in flagella of round spermatids in *Tctex1d2*^+/+^ mice. In elongated spermatids, abnormal expression was observed in *Tctex1d2*^−/−^ mice (top: loss of signal, bottom: ectopic expression). (B) Expression analysis of WDR34 in *Tctex1d2*^−/−^ mice using ICC (green: WDR34, red: acetylated-α-tubulin, blue: hoechst33342). There was no expression of WDR34 was in flagella of round spermatids, as seen with WDR60. In elongated spermatids, abnormal expression was observed in *Tctex1d2*^−/−^ mice (top: loss of signal, bottom: ectopic expression). (C) Expression analysis of DYNLT1 in *Tctex1d2*^−/−^ mice using ICC (green: DYNLT1, red: acetylated-α-tubulin, blue: hoechst33342). The signal was decreased in round spermatids and absent in elongated spermatids in *Tctex1d2*^−/−^ mice. RST, round spermatid; EST, elongated spermatid. (D) Schematic diagram of the defect of cytoplasmic dynein 2 subunits in *Tctex1d2*^−/−^ mice. In*Tctex1d2*^−/−^ mice elongated spermatids, DYNC2LI1, WDR60, WDR34 and DYNLT1 showed abnormal localization and were not assembled in the cytoplasmic dynein 2 complex.

During flagellum formation, the components that constitute the flagella are transported along microtubules. The IFT system is responsible for this transport and consists mainly of IFT proteins, kinesin 2, cytoplasmic dynein 2, and the BBSome complex ^9,37–39^. As *Tctex1d2*^−/−^ mice showed abnormal cytoplasmic dynein 2 localization, we analyzed the expression patterns of other IFT systems. IFT88, an IFT protein, and DYNC2LI1 showed decreased signals and ectopic expression in the flagella of elongated spermatids in *Tctex1d2*^−/−^ mice (fig, S4A, B). In contrast, IFT140 (an IFT protein), KIF3A (a component of kinesin 2), and DYNC2H1 were localized normally in the flagella of elongated spermatids in *Tctex1d2*^−/−^ mice (fig. S4C~E). These results suggest that the IFT system is partially defective in *Tctex1d2*^−/−^ mice. These results are represented in the schematic diagram (fig. 6D). In *Tctex1d2*^+/+^ mice elongated spermatids, the 11 subunits of cytoplasmic dynein 2 form complexes and move on microtubules (fig. 6D). However, *Tctex1d2*^−/−^ mice elongated spermatids, DYNC2LI1, WDR60, WDR34 and DYNLT1 among the 11 subunits showed abnormal localization and were not assembled in the cytoplasmic dynein 2 complex (fig. 6D). Therefore, TCTEX1D2 plays a role in the assembly of cytoplasmic dynein 2 subunits.

## Discussion

In this study, we identified TCTEX1D2 as a dynein light chain important for sperm flagellum formation in mice. *Tctex1d2*^−/−^ mice showed abnormal flagella in round spermatids and disorganization of axonemal components in the flagella. TCTEX1D2 was identified as a component of cytoplasmic dynein 2, based on its interaction with WDR34, WDR60, and DYNLT1, and the expression patterns of these proteins was found to be aberrant in the elongated spermatids of *Tctex1d2*^−/−^ mice. In addition, IFT88 and DYNC2LI1 showed similar abnormal expression patterns in the elongated spermatids of *Tctex1d2*^−/−^ mice. These results suggested that the assembly of cytoplasmic dynein 2 was impaired in *Tctex1d2*^−/−^ mice, consequently affecting the stability of the IFT system. Cilia formation is impaired when the functions of the cytoplasmic dynein 2 and IFT proteins are completely lost ^40–43^. However, the formation of primary cilia is normal in TCTEX1D2 knockout somatic cells in mammalian ^30^, and the formation of motile cilia was normal in *Tctex1d2*^−/−^ mice generated in this study. The expression patterns of cytoplasmic dynein 2, kinesin 2, and IFT proteins were normal in *Tctex1d2*^−/−^ mice (fig. S5A–C). Therefore, the loss of TCTEX1D2 had no significant effect on the IFT system, and cytoplasmic dynein 2 and IFT proteins functioned normally in the sperm flagella and cilia of *Tctex1d2*^−/−^ mice. Indeed, as some components of the IFT system, such as KIF3A, DYNC2H1, and IFT140, are expressed normally in the flagella of *Tctex1d2*^−/−^ mice, it can be concluded that the IFT system works properly during sperm flagellum and cilia formation in *Tctex1d2*^−/−^ mice. Moreover, TCTEX1D2 was expressed in the flagella of round spermatids, and *Tctex1d2*^−/−^ mice showed abnormal flagella in early round spermatids, whereas WDR34 and WDR60 were not expressed in the flagella of round spermatids. As flagellar abnormalities in *Tctex1d2*^−/−^ mice occurred before WDR34 and WDR60 expression, TCTEX1D2 has some functions prior to its function as cytoplasmic dynein 2. Considering these points, the impairment in cytoplasmic dynein 2 by *Tctex1d2* deficit has only a minor effect on sperm flagellum formation therefore, it should be other major causes the flagellar disorders seen in *Tctex1d2*^−/−^ mice.

Another explanation is that TCTEX1D2 functions as an axonemal structure. In *Chlamydomonas*, Tctex2b (a homolog of mouse TCTEX1D2) is present in the axoneme as the light chain of the inner dynein arm ^32,44^. In the present study, we found that TCTEX1D2 was localized in the sperm axoneme and interacted with WDR63 and WDR78. Thus, it was confirmed that TCTEX1D2 is present in the axoneme as a light chain of the inner dynein arm. Transmission electron microscopy revealed that the inner dynein arm was aberrant in the axoneme of the sperm of *Tctex1d2*^−/−^ mice. These results indicate that TCTEX1D2 is important for inner dynein arm assembly. In a previous study, it was shown that *Wdr63* knockout mice exhibited deficiency of sperm flagella and a disruption of the axoneme structure similar to what was seen in *Tctex1d2*^−/−^ mice ^45^. Disorganization and structural instability of the inner dynein arm causes sperm flagellar dysplasia. Therefore, flagellar disorders observed in *Tctex1d2*^−/−^ mice are mainly owing to defects in the inner dynein arm. We intended to perform ICC studies using anti-WDR63 and anti-WDR78 antibodies in *Tctex1d2*^−/−^ mice. However, we could not confirm whether the loss of TCTEX1D2 affects the dynamics of WDR63 and WDR78.

Despite *Tctex1d2*^−/−^ mice having a severely phenotype in sperm flagella, there were no defects in the cilia of the ventricle, trachea, and oviduct. This difference is likely owing to the fact that TCTEX1D2 expressed in the cilia of the ventricles, trachea and oviduct functions only as cytoplasmic dynein 2, not as inner dynein arm, thus weakening the effects of TCTEX1D2 deficit. In a previous study, it was shown that the axonemal components and proteins necessary for axoneme formation differed between sperm flagella and cilia ^16^. Therefore, the function of TCTEX1D2 as an inner dynein arm is unique to sperm flagella. On the other hand, sperm flagella were not completely absent in *Tctex1d2*^−/−^ mice, normally formed up to a certain length in the steps 2-3 round spermatids. As flagella are characterized by a much greater length than cilia, flagellum formation may be undergone a specific change to retain TCTEX1D2 inside the axonemes to maintain the length of flagella. A phenotype similar to that observed in the sperm of *Tctex1d2*^−/−^ mice was observed with CFAP61 deletion. CFAP61, which is a conserved component of the calmodulin- and radial spoke–associated complex of cilia and flagella ^46,47^, when deleted in mice, resulted in motile cilia of normal length; however, the sperm flagella failed to maintain their length ^47^. Furthermore, some knockout mice with axonemal components show normal formation of motile cilia but severe defects in sperm flagella ^16^. Therefore, despite the sperm flagella being regarded as having a structure similar to that of motile cilia, unique molecular mechanisms give rise to the characteristic length and internal structure during its formation.

In the dynein light chain related to cilia and flagella formation, *Tctex1d2* was the first gene to be identified as leading to severe defects in sperm flagella when knocked out in mice. This study showed that dynein light chains, like other chains, play an essential role in sperm flagellum formation in mammals. During cilia formation in mammalian somatic cells, it is known that TCTEX1D2 functions as cytoplasmic dynein 2 ^11,18^. However, during sperm flagellum formation, TCTEX1D2 functions as an inner dynein arm in addition to its function as cytoplasmic dynein 2. Thus, dynein light chains are involved in many processes related to sperm flagella from formation to structural maintenance in mammals. *Tctex1d2*^−/−^ mice showed defective sperm flagella, whereas the Tctex2b mutant *Chlamydomonas* did not show defective flagella but only reduced motility ^32,44^. This indicates that the function of homologous genes differs among species, and it is important to analyze each species.

Multiple morphological abnormalities of the sperm flagella (MMAF) is a male infertility with genetic component, one of the most severe form of sperm defect ^48^. MMAF is characterized abnormal flagella, including short, coiled, absent and of irregular caliber flagellar ^49^. A number of genes have been identified to be associated with MMAF ^50^, yet only 35-60% of MMAF cases have been identified and need for further study to discover new MMAF-related genes ^51,52^. The phenotype of *Tctex1d2*^−/−^ mice is similar to cases seen in MMAF, suggesting that Tctex1d2 deficiency may cause MMAF in human. WDR63, which interacted with TCTEX1D2, is an MMAF-related gene and biallelic variants in WDR63 were identified in infertility men ^45^. Therefore, it is possible that TCTEX1D2 is also a candidate gene causing MMAF. Our study leads to uncover the new MMAF causative genes and could be used as a genetic diagnostic indicator for male infertility.

In summary, we identified *Tctex1d2* as a novel gene essential for the formation of mouse sperm flagella. *Tctex1d2*^−/−^ mice exhibited a severely phenotype only in the sperm flagella, whereas motile cilia had normal structure and function. TCTEX1D2 is weakly involved in the assembly of cytoplasmic dynein 2 and IFT proteins and is important for the assembly and structural stability of the inner dynein arm; this function is unique to sperm flagellum formation, and is not a part of cilia formation. These findings provide novel insights into the function of the dynein light chain in sperm flagellum formation and contribute to the elucidation of the differences between the formation mechanism of sperm flagella and cilia.

## Materials and Methods

### Animals

All mice were C57BL/6N background. C57BL/6N mice were purchased from SLC and maintained under a 12-h light/dark cycle. All animal care and experimental procedures were conducted according to protocols approved by the Tohoku University Institutional Animal Care and Use Committee (2019noudou-003-02, 2019noukumikae-030-06).

### Generation of Tctex1d2^−/−^ mice and Tctex1d2 - 3×FLAG mice

*Tctex1d2* knockouts of the C57BL/6N strain were generated using *i* - GONAD as previously reported ^53^. Briefly, crRNA was designed with the following sequences: 5’-GGTTTGAACCACTACGGTGT-3’ and 5’-TATTCAAGAACTAAGCCCTC-3’. In addition, crRNA and tracrRNA (Integrated DNA Technologies) were annealed at 94 °C for 2 min, mixed with Cas9 nuclease, and injected into the oviduct at E0.7 of plug confirmed mice. Primer sequences for genotyping are given below.

Fw1:5’-ATGCGTGCAAGGATGAGCTA-3’, Rv1:5’-AGGATTTGATCCCCAGCGTG-3’, and Rv2: GCTGGACAACCAGGAAGTGT.

The PCR product from the mutant gDNA was sequenced by Sanger sequencing to confirm successful deletion. To fix the mutation in the strain, the F0 founder was mated with wild-type (WT) mice, and the resulting F1 heterozygous male and female mice were crossed to obtain an F2 generation containing homozygous mutants. The established strain was maintained by sibling breeding and used in the current study.

The method for the generating *Tctex1d2* - 3×FLAG mice was the same as described above. crRNA was designed with the following sequence: 5’-TTGGACCAGACGCAGGGATA-3’. ssODN was designed with the following sequence: 5’-GAGCCGCAACCGACCACACTTCCTTCTAGAGATCTGTTGGACCAGCCGCTGAGCTATGGACTACAAGGACCACGACGGCGATTATAAGGATCACGACATCGACTACAAAGACGACGATGACAAGGCTGTGTCCTTCCGCGGCCTGTCCTTATCGGCTCACTCCGAGGGCTTGAGTGAGGTCG-3’. crRNA and tracrRNA (Integrated DNA Technologies) were annealed, mixed with ssODN and Cas9 nuclease, and injected into the oviduct.

### Fertility test

A single 12-week-old male mouse (*Tctex1d2*^+/+^ or *Tctex1d2*^−/−^) was caged with two 8-week-old female WT mice for 4 weeks. Mating was verified by monitoring the presence of vaginal plugs. The plugged females were then replaced with fresh mice. Litter size was scored for the females that had plugs.

### Histological analysis

Mice were euthanasia, and the testes, epididymis, brain, trachea, and oviducts were collected from *Tctex1d2*^+/+^ and *Tctex1d2*^−/−^ mice. Testes and epididymides were fixed in Bouin’s solution, whereas brain, trachea, and oviducts were fixed in 4 % paraformaldehyde. The fixed tissues were gradually dehydrated using graded series of ethanol and xylene at different concentrations and finally embedded in paraffin. Sections were rehydrated, the testes and epididymis were stained with periodic acid–Schiff (PAS), and the ventricle, trachea, and oviduct were stained with hematoxylin-eosin and examined under a microscope (Olympus BX50).

### Immunohistochemistry

Immunohistochemistry (IHC) studies were performed as previously described ^54^. Tissue sections were placed in antigen retrieval buffer (pH 6.0; citrate buffer) for 20 min at 90 °C. The following primary antibodies were used: rabbit anti-DYKDDDDK tag antibody (20543-1-AP, Proteintech), mouse anti-acetylated α tubulin antibody (sc-23950, Santa Cruz), rabbit anti-TCTEX1 antibody (ab202583, Abcam), rabbit anti-IFT88 antibody (13967-1-AP, Proteintech), and rabbit anti-KIF3A (13930-1-AP, Proteintech). The samples were examined under a fluorescence microscope (Keyence BZ-X710).

### Immunocytochemistry of testicular germ cells

For staining individual cells, samples were prepared using the following protocol. Seminiferous tubules were fixed with 2 % paraformaldehyde containing 0.1 % Triton X-100 for 10 min at room temperature (RT). Pieces of tubules were placed in a drop of the fixing solution on a glass slide, and the pieces were minced in the drop. The slides were gently tapped on the coverslips using tweezers and frozen in liquid nitrogen for 30 s and coverslips were removed. The slides were washed thrice with PBS for 5 min and immunostained. The slides were treated with blocking buffer (Blocking One, Nacalai Tesque, 03953-95) for 1 h at RT and incubated with primary antibody overnight at 4 °C. The following primary antibodies were used: mouse anti-acetylated α tubulin antibody (sc-23950, Santa Cruz), mouse anti-FLAG M2 antibody (F1804, Sigma-Aldrich), rabbit anti-WDR60 antibody (NBP1-90437, Novus Biologicals), rabbit anti-WDR34 antibody (NBP1-88805, Novus Biologicals), rabbit anti-TCTEX1 antibody (ab202583, Abcam), rabbit anti-DYNC2H1 antibody (MBS7050711, MyBioSource), rabbit anti-DYNC2LI1 antibody (PA5-98717, Thermo Fisher Scientific), rabbit anti-IFT88 antibody (13967-1-AP, Proteintech), rabbit anti-KIF3A (13930-1-AP, Proteintech), and rabbit anti-IFT140 antibody (17460-1-AP, Proteintech). Acrosomes were stained with Alexa Fluor 568–conjugated Lectin PNA (L32458; Thermo Fisher Scientific). The samples were examined under a fluorescence microscope (Keyence BZ-X710).

### Western blotting

The tissues (brain, trachea, lung, heart, liver. spleen, kidney, testis) were lysed using RIPA buffer (Nacalai Tesque, 08714-04) containing 1 % phosphatase inhibitor (Nacalai Tesque, 07575-51), and incubated for 30 min on ice. The lysates were centrifuged at 15,000 × *g* for 15 min at 4 °C and eluted with sample buffer for 5 min at 100 °C. The samples were separated by SDS-PAGE, transferred to a PVDF membrane, and treated with a commercial blocking buffer (Blocking One, Nacalai Tesque, 03953-95) for 1 h at RT. The membranes were then incubated with primary antibody overnight at 4 °C. The following primary antibodies were used: rabbit anti-TCTEX1D2 antibody (PA5-116015, Thermo Fisher Scientific) and mouse anti-β-actin antibody (Santa Cruz sc-47778). After washing three times with TBS-T for 5 min, the membranes were incubated with anti-rabbit or anti-mouse HRP-conjugated antibodies (W4011 and W4021, Promega) for 1 h at RT. Images were captured using an ImageQuant LAS 500 (Cytiva).

### Immunoprecipitation

Testes were lysed using RIPA buffer (Nacalai Tesque, 08714-04), and incubated for 30 min on ice. The lysates were centrifuged at 15,000 × *g* for 15 min at 4 °C and were incubated for overnight at 4 °C with anti-FLAG M2 antibody (F1804, Sigma-Aldrich). Subsequently, the lysates were conjugated with 25 µl Protein G Mag Sepharose™ (28944008, cytiva) for 2 h at 4 °C. The immune complexes were washed three times with TBS-T for 5 min and eluted with sample buffer for 10 min at 100 °C. Samples were separated using SDS-PAGE and western blot analysis was subsequently performed. Membranes were stripped with a stripping buffer (Nacalai Tesque, 05364-55), and used for other antibody reactions, as required. The following primary antibodies were used for western blotting: rabbit anti-TCTEX1D2 (PA5-116015, Thermo Fisher Scientific), rabbit anti-WDR60 (NBP1-90437, Novus Biologicals), rabbit anti-WDR34 (NBP1-88805, Novus Biologicals), rabbit anti-TCTEX1 (ab202583, Abcam), mouse anti-WDR78 (sc-390633, Santa Cruz Biotechnology), and rabbit anti-WDR63 antibody (NBP2-32639, Novus).

Images were captured using the ImageQuant LAS 500 (cytiva).

### Fractionation of spermatozoa

Cauda epididymis spermatozoa were suspended in 1 % Triton X-100 lysis buffer (50 mM NaCl, 20 mM Tris-HCl, pH 7.5, protease inhibitor mixture) and incubated for 2 h at 4 °C. The sample was centrifuged at 15,000 × *g* for 10 min to separate the Triton-soluble fraction (supernatant) from the Triton-resistant fraction (pellet). The pellet was resuspended in 1 % SDS lysis buffer (75 mM NaCl, 24 mM EDTA, pH 6.0, protease inhibitor mixture) and incubated for 1 h at RT. The sample was centrifuged at 15,000 × *g* for 10 min to separate the SDS-soluble fraction (supernatant) and the SDS-resistant fraction (pellet). The pellet was dissolved in sample buffer for 10 min at 100 °C. Samples were separated using SDS-PAGE and western blot analysis was subsequently performed. The following primary antibodies were used: rabbit anti-DYKDDDDK tag antibody (20543-1-AP, Proteintech), rabbit anti-GLUT3 antibody (20403-1-AP, Proteintech), mouse anti-acetylated α tubulin antibody (sc-23950, Santa Cruz), and rabbit anti-AKAP3 antibody (13907-1-A, Proteintech). The samples were examined under a fluorescence microscope (Keyence BZ-X710).

### Immunofluorescence for spermatozoa

Cauda epididymis spermatozoa was incubated in human tubular fluid (HTF) medium for 1 h at 37.5 °C with 5 % CO_2_, and swim-up spermatozoa was collected and washed with PBS. The suspension was fixed in 4 % paraformaldehyde for 10 min at RT and air dried on slide glass. The slides were permeabilized with 0.2 % Triton X-100 for 10 min at 37 °C and were washed three times in PBS at 5 min. The slides were treated with blocking buffer (Blocking One, Nacalai Tesque, 03953-95) for 1 h at RT and incubated with primary antibody overnight at 4 °C. The following primary antibodies were used: rabbit anti-DYKDDDDK tag antibody (20543-1-AP, Proteintech) and mouse anti-acetylated α tubulin antibody (sc-23950, Santa Cruz). The samples were examined under a fluorescence microscope (Keyence BZ-X710).

### Sperm motility, count, and abnormal sperm ratio analyses

Cauda epididymis sperm was incubated in HTF sodium for 1 h at 37.5 °C with 5 % CO_2_. Sperm motility parameters were evaluated using a computer-assisted sperm analysis (CASA) system (SMAS; DITECT). Sperm count was analyzed using a hemocytometer. Abnormal sperm ratios were analyzed using phase-contrast microscopy, and 200 sperms from each sample were observed.

### Transmission electron microscopy

*Tctex1d2*^+/+^ or *Tctex1d2*^−/−^ mice testes were cut into 1-mm-thick sections and were fixed with 2.5 % glutaraldehyde in 0.1 M phosphate buffer (PB) for 1.5 h at RT. After the tissues were washed in PB three times, they were additionally fixed with 1 % OsO_4_ in 0.1 M PB for 1.5 h at 4 °C. The tissues were dehydrated using a graded ethanol series for 10 min at RT, which was then replaced with 50 % propylene oxide (PO) in ethanol followed by 100 % PO for 10 min each at RT. The tissues were permeabilized with a mixture of PO and epoxy resin at RT (PO: epoxy resin = 2:1 for 2 h, PO: epoxy resin = 1:1 for 4 h, PO: epoxy resin = 1:2 for 12 h, only epoxy resin for 24 h). The tissues were embedded in epoxy resin for 12 h at 37 °C, for 12 h at 45 °C, and for 12 h at 60 °C. Ultrathin sectioning, staining, and imaging were performed at an external lab.

### Statistical analyses

The unpaired Student’s *t*-test (Microsoft Office Excel; Microsoft Corporation) was used for comparisons with *Tctex1d2*^+/+^ mice. Values are represented as the mean ± SE. Differences were considered significant at *p* < 0.05, indicated as *p* < 0.05 (*).

## Author contributions

Ryua Harima, Conceptualization, Methodology, Validation, Formal analysis, Investigation, Resources, Data curation, Writing – original draft preparation, Writing – review & editing, Visualization, Project administration, Funding acquisition; Kenshiro Hara, Resources, Writing – original draft preparation, Writing – review & editing, Funding acquisition; Kentaro Tanemura; Resources, Supervision, Writing – original draft preparation, Writing – review & editing, Funding acquisition.

## Acknowledgments

We would like to thank Editage (www.editage.com) for English language editing.

## Supplementary Material

**Supplement figure. 1.**
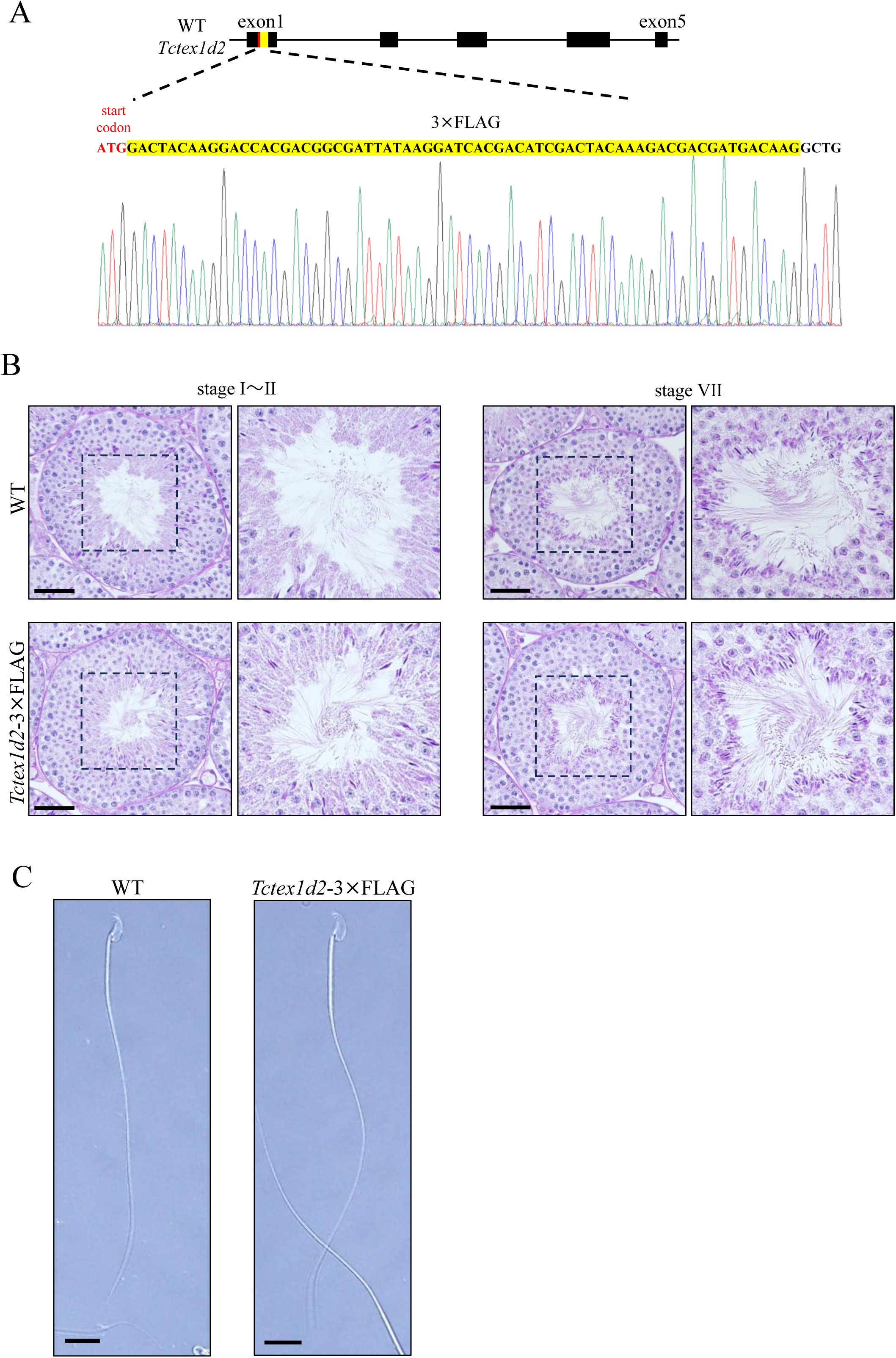
Generation of *Tctex1d2* - 3×FLAG mice. (A) Schematic representation of *Tctex1d2* - 3×FLAG mice and the results of Sanger sequencing. 3×FLAG was knocked immediately after the start codon. Sanger sequencing indicated that the 3 × FLAG nucleotide sequence was knocked-in within the target region. (B) PAS-H staining of seminiferous tubules of WT and *Tctex1d2* - 3×FLAG mice. Scale bars are 50 µm. (C) Phase contrast microscopy of cauda epididymis sperm. There were no differences of sperm morphology between WT and *Tctex1d2* - 3×FLAG mice. Scale bars are 10 µm.

**Supplement figure. 2.**
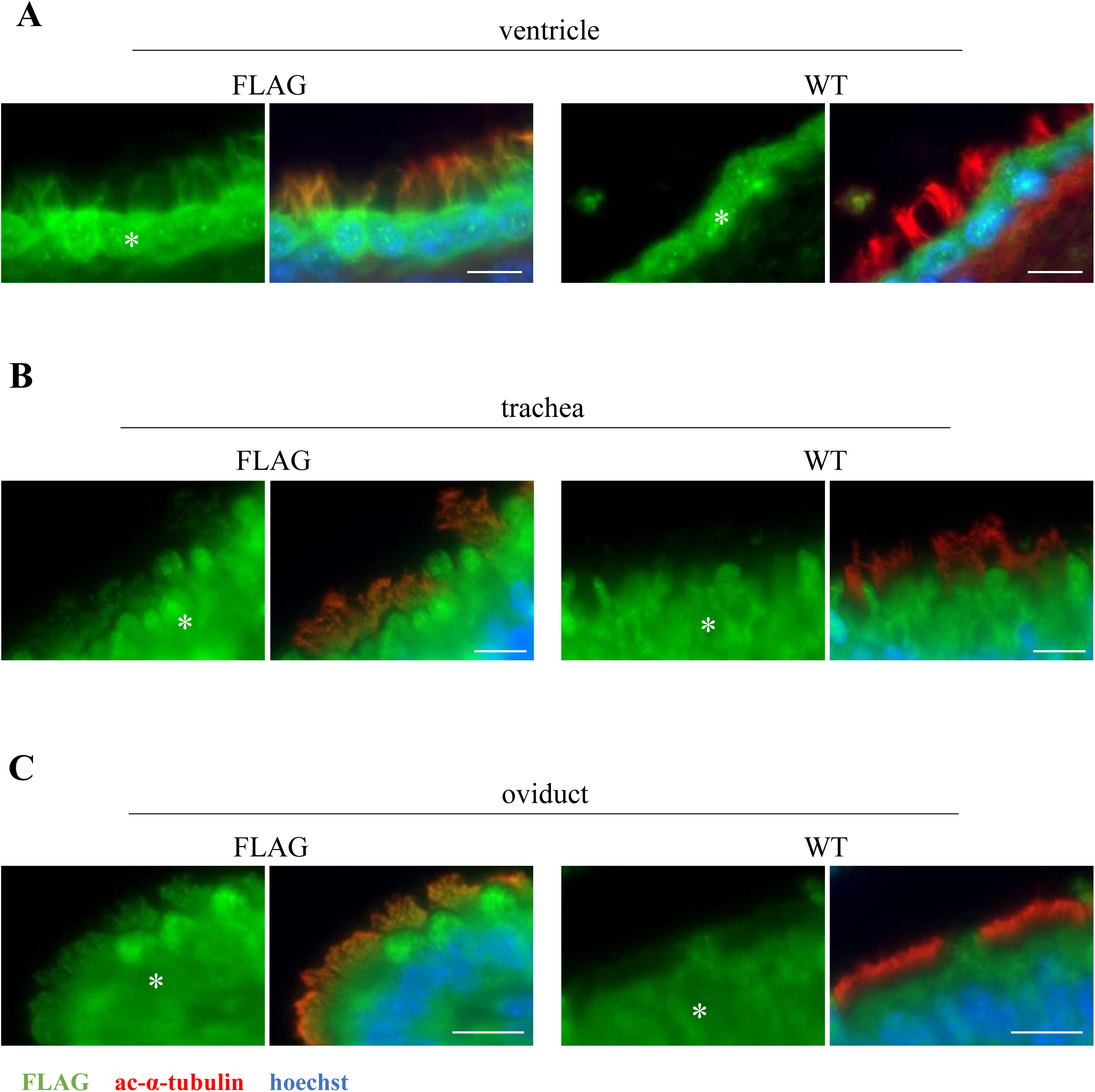
TCTEX1D2 was also localized to motile cilia. IHC of WT and *Tctex1d2* - 3×FLAG in tissues with motile cilia to analyze the expression of TCTEX1D2 (green: anti-FLAG, red: anti-acetylated-α-tubulin, blue: hoechst33342). (A) ventricle, (B) trachea, and (C) oviduct. TCTEX1D2 is expressed in the cilia of the ventricle, trachea, and oviduct. (*); The signals found in the epithelial cells of each tissue are non-specific. Scale bars are 10 µm.

**Supplement figure. 3.**
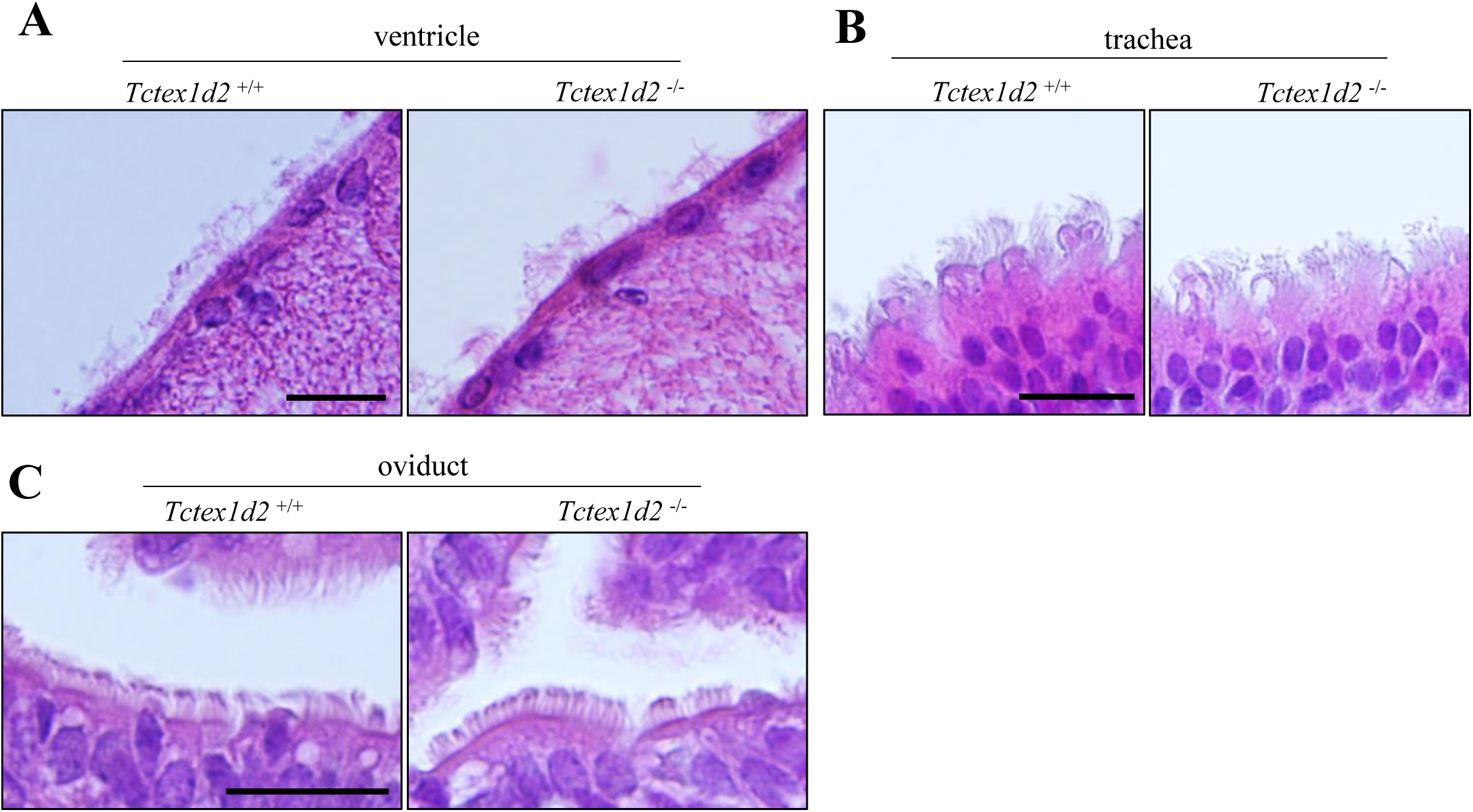
Motile cilia in *Tctex1d2*^−/−^ mice were normal. H&E stain of tissues with motile cilia. (A) ventricle, (B) trachea, and (C) oviduct. In motile cilia, there were no differences in morphology between *Tctex1d2*^+/+^ and *Tctex1d2*^−/−^ mice. Scale bars are 20 µm.

**Supplement figure. 4.**
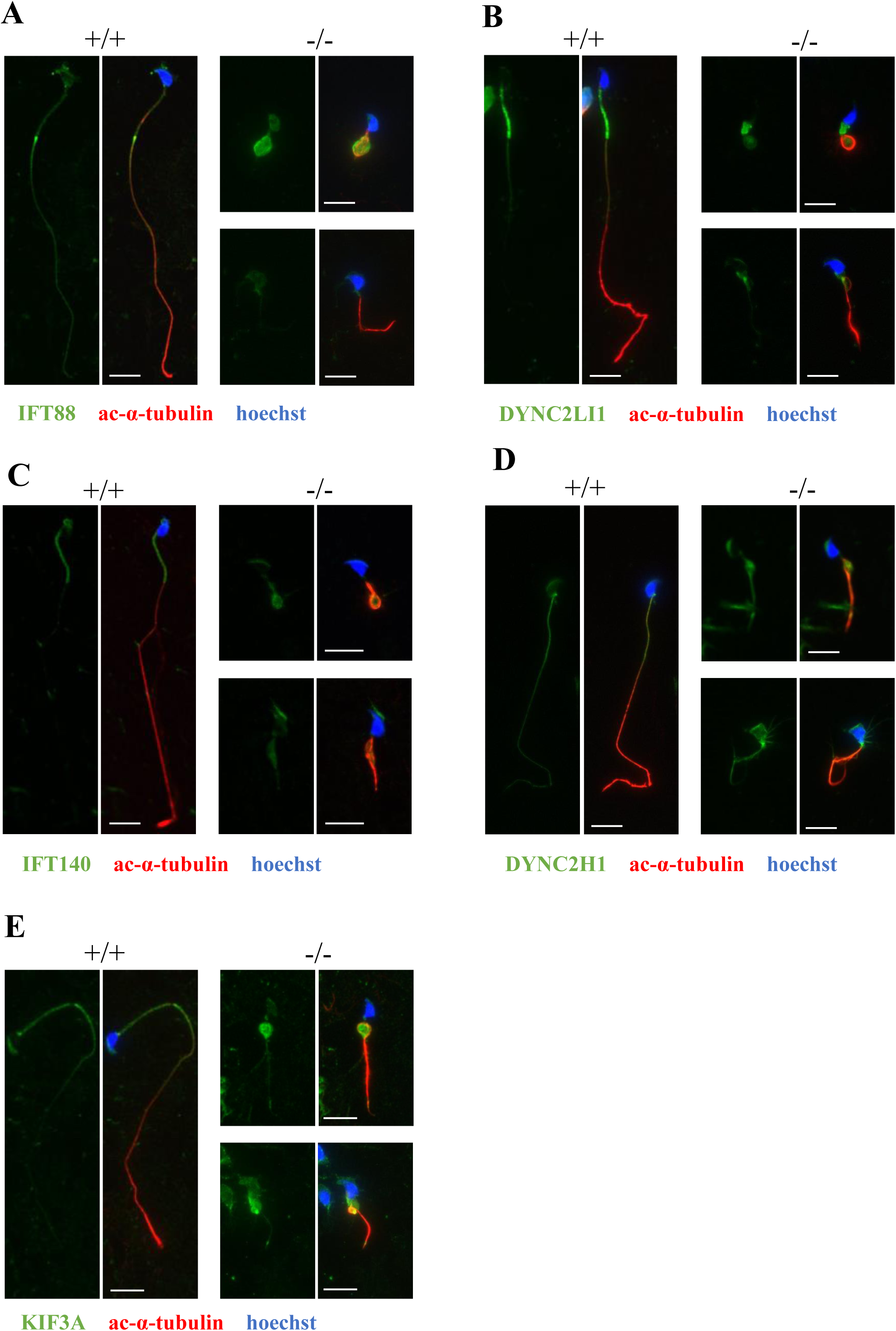
*Tctex1d2*^−/−^ mice exhibited abnormal protein localization in the IFT system. Expression analysis of cytoplasmic dynein 2 (DYNC2H1 and DYNC2LI1), kinesin 2 (KIF3A), and IFT proteins (IFT88, 140) in elongated spermatids of *Tctex1d2*^−/−^ mice using ICC (red: anti-acetylated-α-tubulin, Blue: hoechst33342). (A) IFT88, (B) DYNC2LI1, (C) IFT140, (D) DYNC2H1, and (E) KIF3A. IFT88 and DYNC2LI1 showed a reduced signal and were ectopically expressed in the flagella of *Tctex1d2*^−/−^ mice, whereas IFT140, DYNC2H1, and KIF3A were expressed normally in *Tctex1d2*^−/−^ mice. Scale bars are 10 µm.

**Supplement figure. 5.**
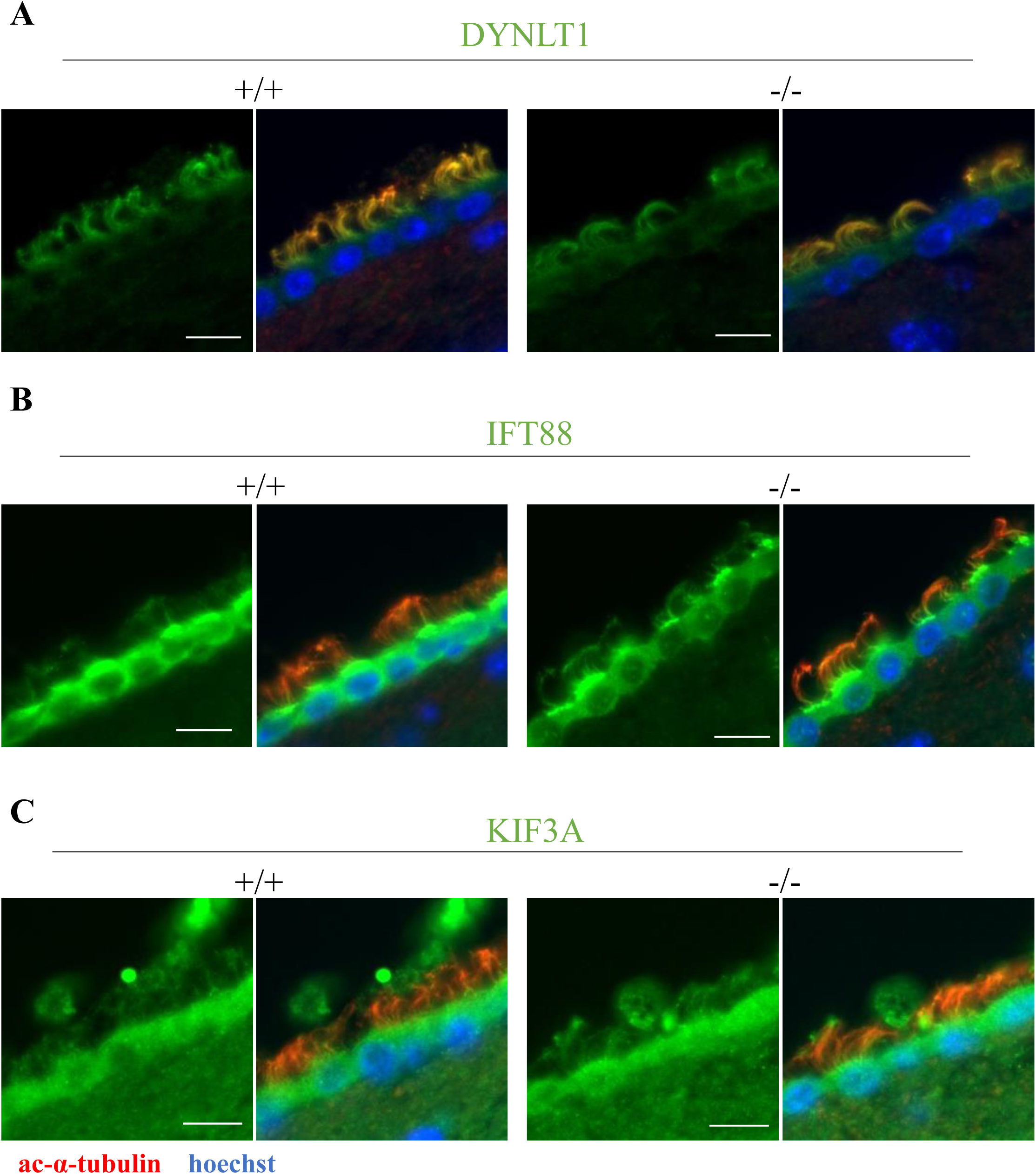
The IFT system was normal in the cilia of the ventricles. Expression analysis of cytoplasmic dynein 2 (DYNLT1) and IFT protein (IFT88) in the ventricle of *Tctex1d2*^−/−^ mice using IHC (red: anti-acetylated-α-tubulin, blue: hoechst33342). In *Tctex1d2*^−/−^ mice, the two proteins showed abnormal expression in the flagella of elongated spermatids, but not in the ventricle. Scale bars are 10 µm.

